# A MAP of tumor-host interactions in glioma at single cell resolution

**DOI:** 10.1101/827758

**Authors:** Francesca Pia Caruso, Luciano Garofano, Fulvio D’Angelo, Kai Yu, Fuchou Tang, Jinzhou Yuan, Jing Zhang, Luigi Cerulo, Davide Bedognetti, Peter A. Sims, Mario Suvà, Xiao-Dong Su, Anna Lasorella, Antonio Iavarone, Michele Ceccarelli

## Abstract

Single-cell RNA sequencing is the reference technique to characterize the heterogeneity of tumor microenvironment and can be efficiently used to discover cross-talk mechanisms between immune cells and cancer cells. We present a novel method, single cell Tumor-Host Interaction tool (`scTHI`), to identify significantly activated ligand-receptor interactions across clusters of cells from single-cell RNA sequencing data. We apply our approach to uncover the ligand-receptor interactions in glioma using six publicly available human glioma datasets encompassing 71 patients. We provide a comprehensive map of the signalling mechanisms between malignant cells and non-malignant cells in glioma uncovering potential novel therapeutic targets.

## INTRODUCTION

The interaction between malignant cells and their microenvironment is critical for both normal tissue homeostasis and tumor growth, as the immune system can eliminate cancer cells that present neoantigens recognized by receptors of the adaptive immune system or express ligands for activating receptors on innate immune cells (Schreiber et al., 2011). The composition of the various cell types making up the microenvironment can significantly affect the way in which the immune system activates cancer rejection mechanisms (Bedognetti et al., 2016; Joyce and Fearon, 2015; Zhang et al., 2019) and influences the response to immune therapies (Ceccarelli et al., 2016; Cristescu et al., 2018). Therefore, the elucidation of the tumor-host interaction mechanisms plays a crucial role in the understanding of the tumor growth and evolution (Angelova et al., 2018; Bedognetti et al., 2019) and for the identification immuno-oncology therapeutic targets (Hoos, 2016). Immune checkpoint inhibitor (ICI) therapies are aimed to targeting the specific cell-cell interaction between PD1 and PD-L1 or CTLA4 and B7-1/B7-2 (Pardoll, 2012). The identification of novel interactions that characterize tumor-types and shape the immune response has also important clinical implications can also help to better stratify patients (Zhang et al., 2019) and predict response to ICI (Roelands et al., 2019).

Single-cell RNA sequencing is the reference technology for the quantification and phenotyping of tumor microenvironment at high resolution (Azizi et al., 2018; Svensson et al., 2018) allowing to measure the composition of individual immune/stromal compartments making up the microenvironment. This technique can be also used for a better elucidation of the tumor-host signalling mechanisms (Kumar et al., 2018) and the identification of tissue-specific interactions at an unprecedented spatially-solved level of details (Govek et al., 2019).

Glioma are characterized by the worst survival among brain tumor malignancies (Ceccarelli et al., 2016). In particular Glioblastoma (grade IV glioma) is the most frequent type of primary brain tumor being an invariably lethal tumor type with median survival below 15 months (Stupp et al., 2005). In glioma, higher mutational load is associated with increased tumor aggressiveness (Zhang et al., 2019). The tumor microenvironment of GBM is dominated by myeloid-derived cells, mostly blood-derived macrophages and resident microglia (Quail and Joyce, 2017; Wang et al., 2018; Yuan et al., 2018), hampering a productive anti-tumor immunity in GBM and excluding T lymphocytes and (Hussain et al., 2006). Therefore, due to this lymphocyte depleted immunosuppressive microenvironment, several multi-cancer studies led to the exclusion of high-grade glioma patients (Weller and Fontana, 1995). The elucidation of the active ligand-receptors (L-R) interactions in the cross-talk between tumor cells and their microenvironment can help to identify the mechanisms that the transformed cells in glioma use to recruit this immunosuppressive microenvironment and to discover novel therapeutic targets.

Here we exploit single cell data developing `scTHI`, a novel algorithm and tool to identify the ligand-receptor pairs that modulate the tumor-microenvironment cross-talk in glioma. `scTHI` is based on the hypothesis that when patterns of interaction are active, they are also simultaneously and highly expressed in homogeneous cell populations. We also model the autocrine and paracrine signalling effects of L-R partners (Ramilowski et al., 2015). Interestingly, by collecting the largest collection of single cells dataset available up to date, we show that unexpected cross-talk partners are highly conserved across different datasets in the majority of the tumor samples. This suggests that shared cross-talk mechanisms exist in glioma. Our results provide a complete map of the active tumor-host interaction pairs in glioma that can be therapeutically exploited to reduce the immunosuppressive action of the microenvironment in brain tumor.

## RESULTS

We present `scTHI` an R package to discover L-R interactions in single cells. There have been few attempts in scoring such pairs that are mainly based on the average expression of the gene pairs across cell populations. One recent approach is reported in (Kumar et al., 2018) where the authors score interactions by calculating the product of mean receptor expression and mean ligand expression in the respective cell types under examination. The significance of the interaction is evaluated through a one-sided Wilcoxon rank-sum between the median interaction score across samples. This idea is similar to the original approach reported by the authors of the CellPhoneDB (Vento-Tormo et al., 2018) where the mean expression of the gene pair is considered with the constraint that only receptors and ligands expressed in more than 10% of the cells in the analyzed cluster are selected. The significance of the interaction is then evaluated using a random permutation of the samples. The scTHI tool inherited some of these ideas with the main difference that we do not use the mean of the expression as it is heavily influenced by drop-out phenomena. Our scTHI score is based on the percentage of cells in two clusters where the expression of the R-L genes is ranked at the top of the expression profile of every cell of the cluster. Due to the limitations in mRNA capture affecting scRNA sequencing, the use of ranked expression values to compare gene pairs is more stable than just averaging gene expression, reducing the false positive rate in interaction pairs detection. Furthermore, the choice to use as score the percentage of cells expressing a top-rank L-R pair gives priority to those pairs that are more uniformly expressed in two clusters of cells. Overall, this allows us to discard interaction pairs for which one of the two members is highly expressed and the other is not, which instead would be detected using a score based on the average of the expression. Moreover, we explicitly model paracrine and autocrine effects. Since we are particularly interested in paracrine effects, our score penalizes L-R pairs where both partners are highly expressed in the same cell. We assess the significance of the score using a bootstrap method similar to (Vento-Tormo et al., 2018) (see Methods).

The basic workflow of scTHI is presented in **Figure 1.** First, we perform a cell-specific identification using the gene set enrichment Mann-Whitney-Wilcoxon Gene Set Test (mww-GST) (Frattini et al., 2018) based on a collection of 295 gene immune and stromal signatures. We adopt mww-GST since it has been proven to perform better than other enrichment analysis in situations of weak and noisy signals, and therefore can be used in the single cell scenarios with the presence of low detection efficiency and drop-out phenomena. The second step of the pipeline is the scoring of the candidate L-R pairs using the procedure described in the Methods section. Briefly, the method computes for each cluster the percentage of cells where the L-R partners are ranked at the top 20% (this is a tunable parameter). The score is the mean between the two percentages and it prioritizes L-R with paracrine activation removing form the score the autocrine effects (see Methods). The significance of the score is computed by a bootstrap generated null distribution obtained by randomly shuffling the input data.

**Figure 1.**
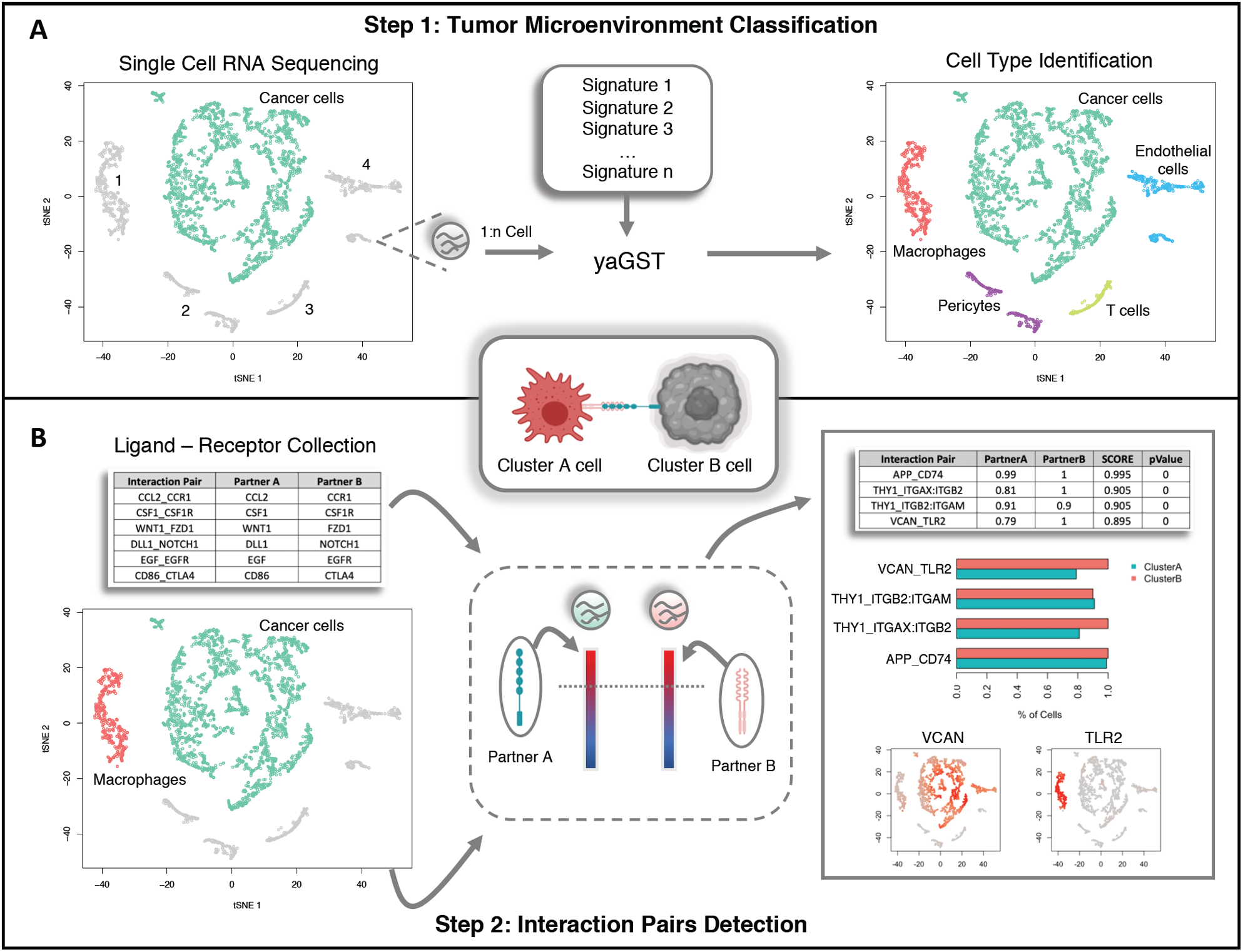
scTHI workflow. Description of scTHI functionalities. (A) In the first step a single sample enrichment analysis based on the Mann-Witney-Wilcoxon Gene Set Test (mww-GST) (38) with a collection of 295 gene immune and stromal signatures is used to nominate the identity of cells of the microenvironment. (B) In the second step, given two clusters of cells, the most significant L-R interactions are identified by assigning a score to every all the 2,548 collected pairs of ligands and receptors.

### scTHI is able to recover validated interactions from single cell data

Many putative ligand/receptor interactions could be identified based on quantitative gene expression evaluation. However, only those occurring among cells spatially close to each other could have a real biological functionality. Goltsev et al. observed that the cellular neighborhood has a profound impact on the expression of protein receptors on immune cells (Goltsev et al., 2018), highlighting that the spatial resolution of infiltrating immune cells and the cancer cells play a key role in defining tumor heterogeneity. Given these assumptions, we evaluated whether `scTHI` is able to detect interactions occurring between cluster of cells spatially close. For this reason, we used the high quality CITE-seq dataset described in Govek et al., where the spatial architecture of murine splenic cells was resolved (Govek et al., 2019).

First, we used `scTHI` to identify all cell types composing the murine spleen described by Govek et al., including T cells (cytotoxic, memory, naïve, regulatory and helper), B cells (follicular, naïve and switched memory), dendritic cells (conventional and plasmacytoid), red pulp macrophages, monocytes-derived macrophages, neutrophils, plasma cells, erythrocytes and erythroid progenitors. We used the signatures in `scTHI` (generated for human) converted to their mouse orthologs. We asked if scTHI was able to detect interactions occurring between clusters of spatially close cells. Govek et al. used CODEX to validate some important ligand/receptor interactions occurring among red-pulp macrophages and monocyte-derived macrophages (C1q-Lrp1), red-pulp macrophages and neutrophils (Hebp1-Fpr2), monocyte-derived macrophages and neutrophils (Anxa1-Fpr1). They identified these interactions on the basis of the proximity of the corresponding cells expressing ligand and receptor in both the CITEseq and CODEX data. interestingly, `scTHI` detected the validated interactions among the top 10 highest scored, without any spatial information, as shown in **Figure S1**. This highlights how our approach, which scored pairs based on rank expression values, is robust and accurate in the identification of relevant L-R interactions.

### Map of non-tumor cells in glioma

Gliomas are primary brain tumors characterized by high levels of intratumor heterogeneity and, despite numerous research advances, the difference in tumor microenvironment composition is still not well understood (Chen and Hambardzumyan, 2018). We collected a single cell glioma dataset integrating six published studies. This allowed us to comprehensively evaluate the composition of the tumor microenvironment, spanning different molecular and histological subtypes of glioma. Overall, we have 45,550 malignant cells and 11,510 non-malignant cells among datasets. We classified all non-malignant cells using scTHI, however, below we report the percentages of specific cell compartments computed using the datasets where the cells did now undergo any gating or selection strategy. Classification of the non-malignant cells (**Table S3**) showed that the most relevant fraction of cells in the glioma microenvironment were myeloid cells (∼57%) divided in macrophages (∼45%) and microglia (∼12%) followed by glial cells (∼19%), vascular cells (∼11%), CD8 T cells (∼4%) and few subpopulations of other cell types including natural killer, neutrophils, dendritic cells, monocytes, mesenchymal stem cells and others (∼9%). As expected, grade IV glioma (GBM) showed the highest percentage of macrophages in their microenvironment (∼52% macrophages and ∼8% microglia) compared to other histological subtypes (Astrocytoma-macrophages = ∼10% and Astrocytoma-microglia = ∼36%; OligoAstro-macrophages = ∼9% and OligoAstro-microglia = ∼21%; Oligodendroglioma-macrophages = ∼1% and Oligodendroglioma-microglia = ∼31%) (**Figure 2**). Interestingly, switching from more aggressive histological phenotypes (i.e. GBM) to less aggressive ones (i.e. Oligodendroglioma) the relative percentage of macrophages decreases, while the percentage of microglia cells increases. These data are in agreement with the hypothesis that gliomas in the early stages of their development primarily contain brain-resident microglia cells, whereas macrophage phenotype is associated with higher grades (Venteicher et al., 2017). Astrocytoma and glioblastoma patients showed also a relevant fraction of vascular cells (∼44% and ∼14%, respectively), probably due to increased microvascular proliferation of these high grade tumors compared to oligodendrogliomas. Regarding the lymphoid populations, T cells represent the most abundant fraction, with a greater number of CD8 cells observed in GBM and Oligo-Astrocytoma.

**Figure 2.**
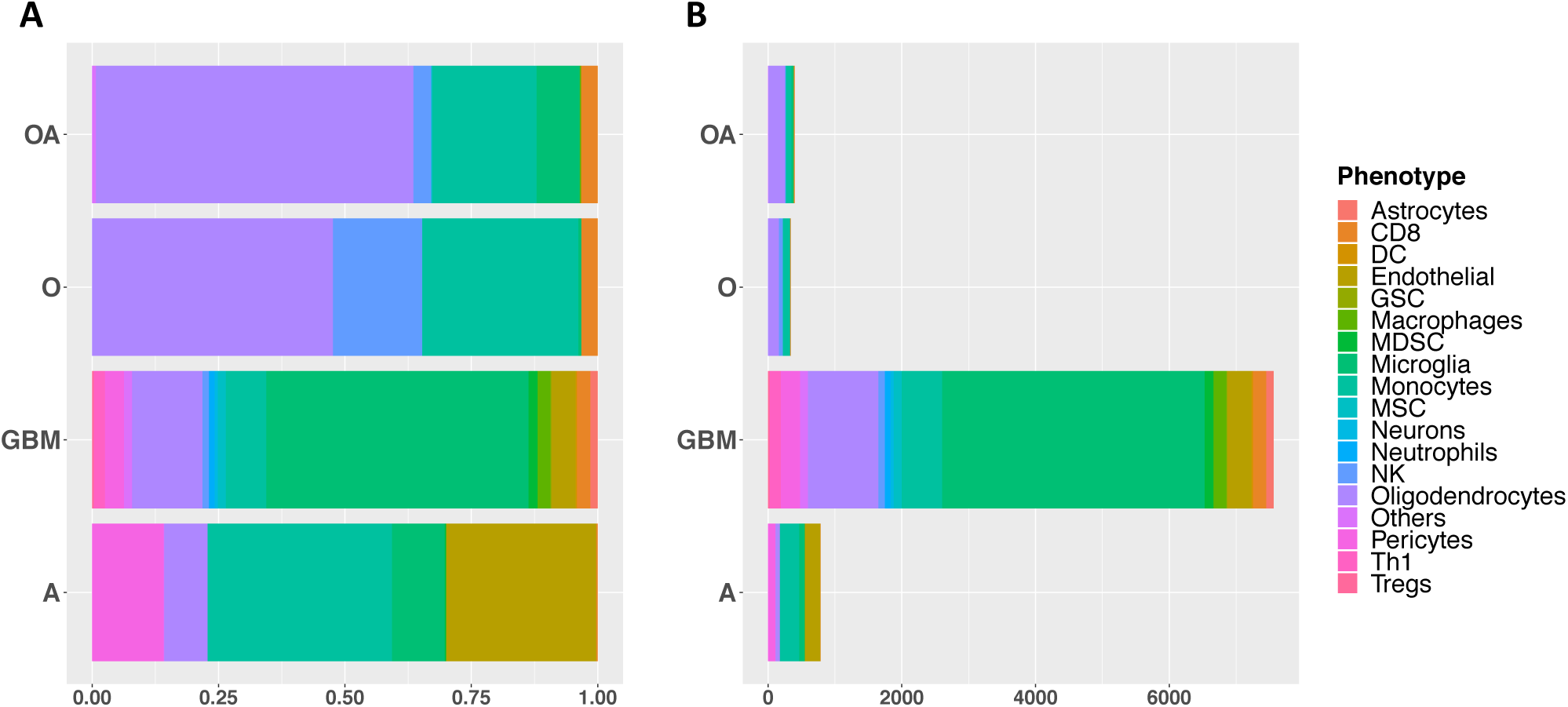
Tumor microenvironment cell type classification in glioma. The barplots show the relative percentage (A) and the number of cells (B) of each cell type identified in the microenvironment of the main histological subtypes of glioma.

We also evaluated whether there is a significant association between the different cell types composing the microenvironment and the molecular glioma subtypes (Wang et al., 2017). We correlated the percentage of cells classified in one of the glioma subtypes with the percentages of non-malignant cell types only for patients in which the cells were not selected with any gate strategy (**Figure S2**). This analysis showed a significant correlation between the Mesenchymal subtype and the presence of macrophages (rho=0.47, p-value=0.015), myeloid-derived suppressor cells (MDSC) (rho=0.56, p-value=0.003), dendritic cells (rho=0.40, p-value=0.039), and astrocytes (rho=0.42, p-value=0.033); Proneural subtypes was significantly associated with the presence of Oligodendrocytes (rho=0.43, p-value=0.028); Classical subtype was significantly associated with the presence of Microglia (rho=0.45, p-value=0.019).

### Cross-talk between mesenchymal GBM tumor cells and myeloid cells in the glioma microenvironment

Since cells of myeloid lineage account for about 50-60% of non-neoplastic cells, we first focused on interactions occurring between tumor and myeloid cells, including bone marrow-derived macrophages and microglia. The analysis was performed on the patients with a sufficient number of detected myeloid cells (n = 39 patients). Each patient was tested to identify both paracrine and autocrine interaction pairs. Only significant L-R pairs were kept, with p-*value* less than 0.05 and the constraint a total scTHI score greater than 0.50 in both clusters. Altogether, we detected 368 significant L-R pairs across datasets, we filtered out all interactions occurring in fewer than 4 patients. About 80% of detected interactions (298 out of 368) showed a considerable autocrine signaling (**Table S4**), the remaining 20% of the identified interactions (n = 70) showed paracrine signaling, where the interaction genes were preferably expressed on only one of the two clusters (**Table S4**). Several of high-scored interactions occurred specifically in few patients due to the typical heterogeneity of the considered tumor. Interestingly, a relevant fraction of detected pairs (n = 56, about 15%) was shared at least 50% or more of patients, suggesting the presence of common tumor-host signaling mechanisms in glioma. Many of the inferred interactions involved genes of chemokine and cytokine family (e.g. CCL5, CCR1, CCRL2), toll-like receptors (e.g. TLR2, TLR4), transforming growth factor genes, tumor necrosis factor genes, MHC proteins (e.g. HLA-E, CD74), growth factors and their receptors (e.g. EGFR, PDGFB, PDGFC, PDGFRA, IGF1, IGF1R), cell adhesion molecules (e.g. integrins), enzymes with inhibitor activity, and metalloproteinases. Gene ontology enrichment analysis revealed that secreted ligands or activated receptors in tumor cells are typically involved in processes of extracellular matrix remodeling, cell chemotaxis, Notch signaling, axonogenesis and gliogenesis (**Figure S3A and Table S5**). Instead, receptors and ligands detected on myeloid cells modulate biological processes such as leukocyte chemotaxis and migration, cell-cell adhesion, cytokine production, myeloid cell differentiation, reactive oxygen species metabolic processes and regulation of vasculature development (**Figure S3B and Table S5**).

The significant interactions identified running scTHI in paracrine mode and most common among all glioma patients, were VCAN-TLR2 (72% of patients) and HBEGF-EGFR (51% of patients) (**Figure 3**). The Versican gene (VCAN) is an extracellular matrix proteoglycan, typically involved in processes of cell adhesion, proliferation, and migration. It is highly expressed in glioma cells where strongly contributes to tumor progression mechanisms (Hu et al., 2015). According to these observations, we found VCAN diffusely expressed in the tumor cells of all the datasets considered (**Figure 4)**. The interaction partner, the Toll-like receptor 2 (TLR2), is a membrane protein expressed on the surface of many cell types, including monocytes and macrophages, and it is involved in pattern recognition signaling pathway and innate immunity activation. TLR2 is highly and specifically expressed only in cells of the microenvironment of myeloid origin, and low expressed in other cell types (**Figure 4B**). The interaction occurring among VCAN and TLR2 represents an effective link between inflammation and tumor progression. Indeed, the Versican protein, released by tumor cells in the extracellular space, binds TLR2 activating multiple cell types in the tumor microenvironment, including myeloid, fibroblasts and endothelial cells, and promotes the production of many proinflammatory cytokines (Kim et al., 2009; Wang et al., 2009). The activation of TLR2 downstream signaling pathway also induces the expression of metalloproteinases involved in extracellular matrix degradation to promote tumor expansion (Hu et al., 2015). The most common interaction with the receptor on the tumor cells was composed by the pair EGFR and HBEGF, that could tend to promote tumor growth. In fact, the Epidermal Growth Factor Receptor (EGFR) is a tumor-promoting receptor commonly amplified in gliomas, and EGF-Like growth factor (HBEGF) is a protein highly expressed in regulatory macrophages with immunosuppressive activity (Edwards et al., 2009). Among other significant interactions of interest identified by scTHI, one involves the macrophage migration inhibitory factor MIF and its receptor CD74 (18% of patients), it plays a role in tumorigenesis, exerting pro-tumorigenic effects such as enhancing proliferation, tumor vascularization and inhibition of apoptosis (Fukaya et al., 2016).

**Figure 3.**
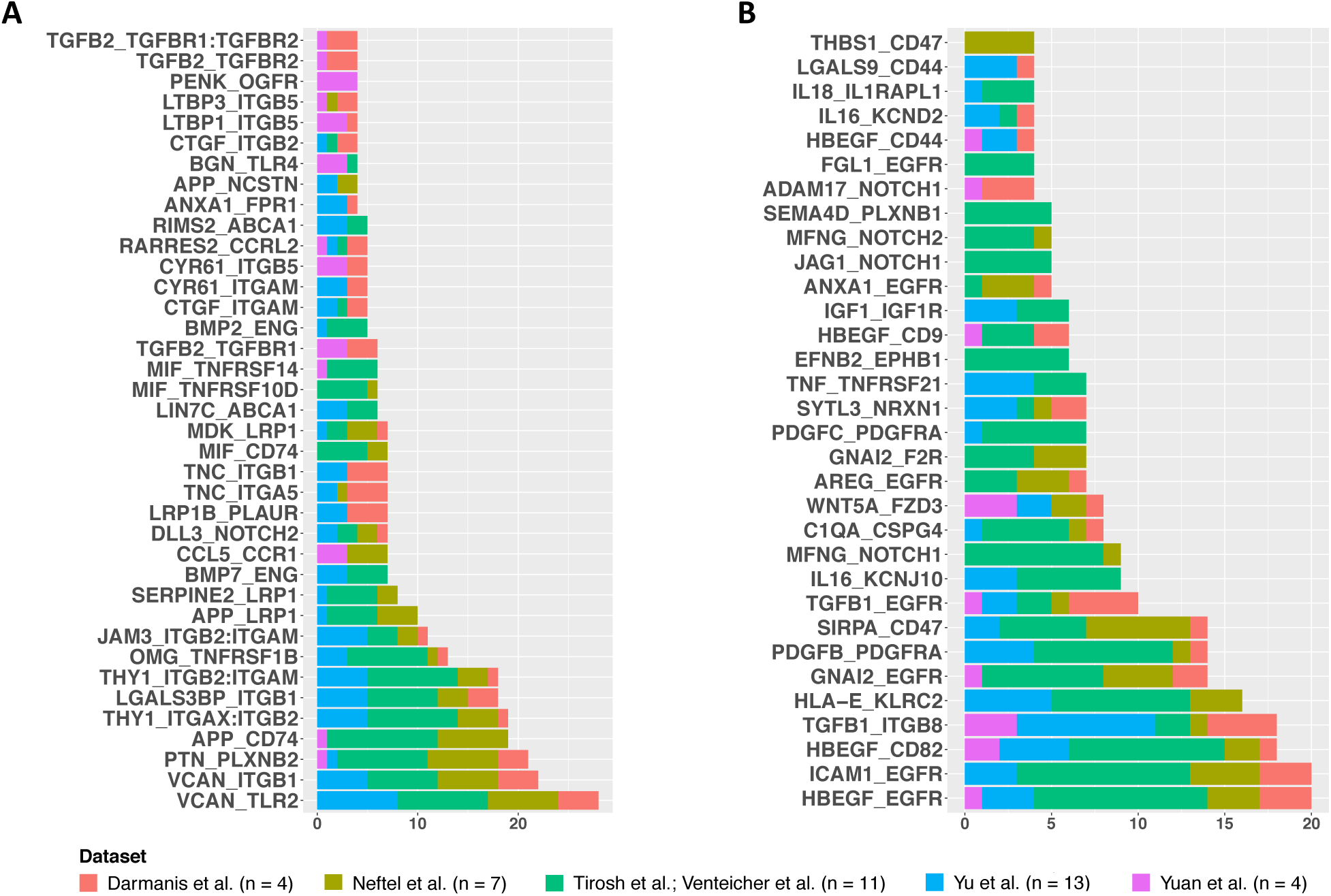
Paracrine tumor-myeloid cell interactions. Barplots show significant paracrine L-R interactions (p-Value <= 0.05 and scTHI score >= 0.50) occurring between tumor and myeloid cells shared in at least 4 patients. On the x axis are shown the number of patients where each interaction occurred. (A) Interaction pairs in which the ligand is expressed on tumor cells and the receptor on myeloid cells. (B) Interaction pairs in which the ligand is expressed on myeloid cells and the receptor on tumor cells.

**Figure 4.**
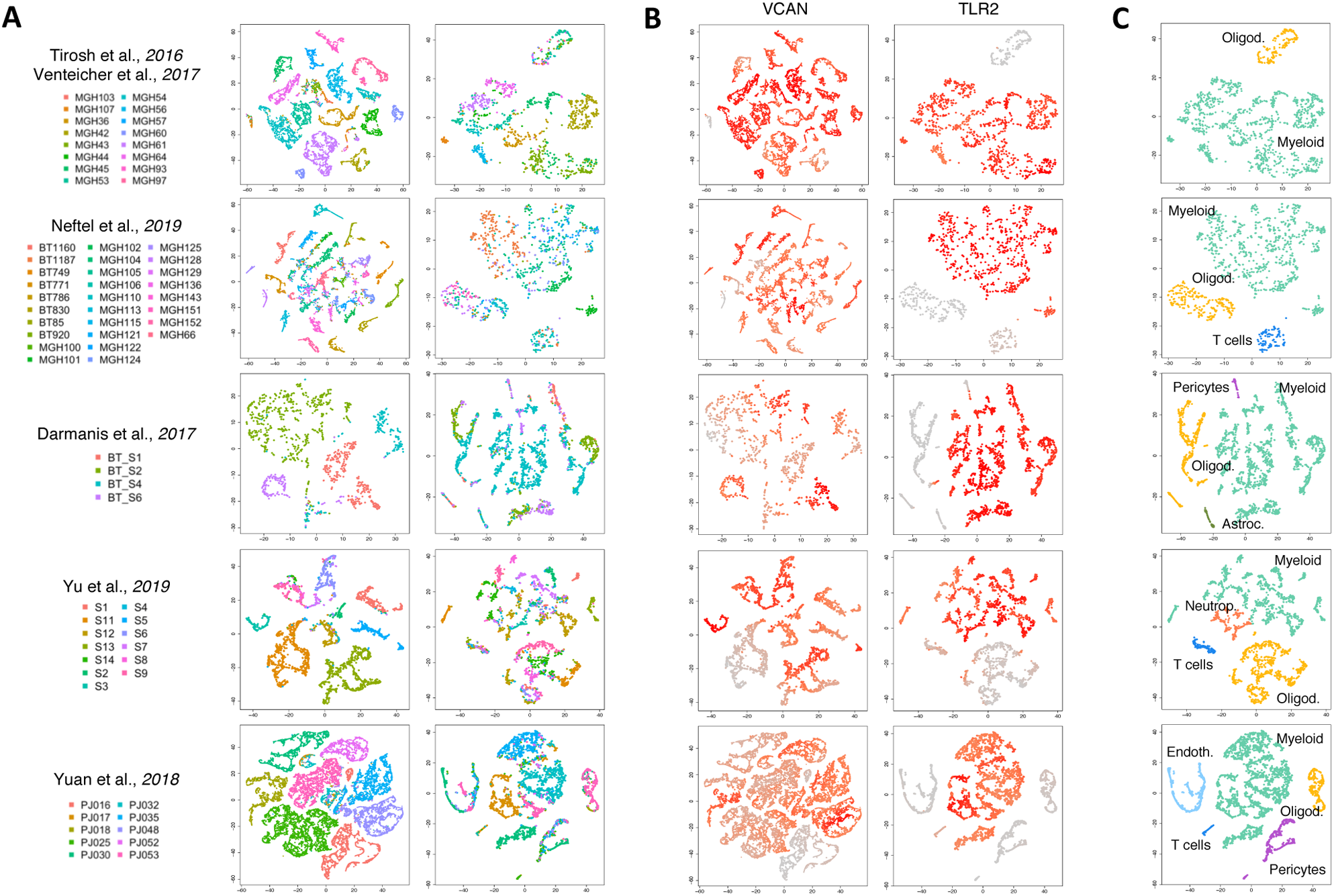
VCAN-TLR2 interaction. TSNE plots of tumor and non-tumor cells for each dataset analyzed. (A) Each cell is colored by patient (on the right non-tumor and on the left tumor cells). (B) Each cell is colored according to expression value of the genes VCAN and TLR2 in tumor and non tumor cells, respectively. (C) tSNE plot of non-tumor cells colored according to cell type classification.

We also found that multiple significant interaction pairs involved ligands (TGFB1, TGFB2), receptors (TGFBR1, TGFBR2) and regulators (LTBP1, LTBP3) of the TGFb signaling pathway. The TGFb pathway is a known driver of immunosuppression in multiple epithelial tumors and in glioblastoma in which it also drives other hallmarks of aggressiveness (cancer stem cells, migration, and invasion,etc.) leading to poor survival (Anido et al., 2010; Kaminska et al., 2013).

The identification of interaction pairs in autocrine mode (**Table S4)** revealed that these signaling are much more conserved among patients than paracrine signaling. As expected, some of the paracrine interactions described above were also identified as autocrine, although they have a preferential paracrine directionality. Among the top scored L-R autocrine pairs, there was an interaction between RPS19 and C5AR1 (**Figure S4**), detected in all the glioma patients tested (n = 39). Although this interaction has never been described in the context of glioma, the Ribosomal protein S19 (RPS19) is up-regulated in breast and ovarian cancers, and its interaction with the C5a receptor 1 (C5AR1), expressed on tumor-infiltrating myeloid cells, has an immunosuppressive effect. RPS19 induces the production of anti-inflammatory cytokines, the activation of Th2 and regulatory T cells, and the reduction of infiltrating CD8 T cells into tumors. It was also noted that reducing RPS19 in tumor cells or blocking the C5aR1-RPS19 interaction decreases RPS19-mediated immunosuppression, impairing the tumor growth (Markiewski et al., 2017). These observations could be translated into gliomas, representing another potential therapeutic target.

### Cross-talk between proneural GBM tumor cells and oligodendrocytes in the glioma microenvironment

Whereas the microenvironment of mesenchymal GBM was massively enriched by myeloid cells, proneural GBM contained a low number of myeloid cells but exhibited significant infiltration by oligodendrocytes, which alone accounted for about 17% of tumor-infiltrating cells. Altogether we tested 30 tumor samples, among the 71, for which we identified an adequate number of oligodendrocytes in their microenvironment. Each individual tumor was tested to identify both paracrine and autocrine interaction pairs, and only significant interactions shared in at least 4 patients were considered. Overall, we found 26 paracrine and 126 autocrine interactions (**Table S6**). Gene ontology enrichment analysis revealed that the cross-talk between tumor cells and oligodendrocytes mainly involve signaling pathways related to cell growth and nervous system development, like as axonogenesis, regulation of neurogenesis, extracellular matrix organization, developmental cell growth, gliogenesis, synapse organization (**Table S7**). Indeed, among the top-scored paracrine interactions (**Figure S5**) there were several ligands and receptors involved in cell-cell adhesion, angiogenesis, and tumorigenesis, such as MDK-LRP2 (12 out of 30 patients) and JAM2-JAM3 (11 out of 30 patients).

The MDK-LRP2, which scored as top interaction among GBM tumor cells (ligand) and oligodendrocyte (receptor) is especially intriguing as overexpression of MDK (midkine) has been shown in several human tumors and recently was reported as driver of aberrant proliferation, poor survival and pharmacological resistance in human glioma (Cheng et al., 2014; Lorente et al., 2011). In many patients, we also detected as significant the interaction occurring between the Platelet Derived Growth Factor Subunit A (PDGFA) gene and its receptor, PDGFRA (11 out of 30 patients). PDGFA is a classical marker of oligodendrocytes required for normal oligodendrocyte development. However, the overexpression of its receptor is a hallmark of proneural GBM, where it plays a critical role during tumor development and progression (Song et al., 2018). We found that ∼72% of cells expressing PDGFRA at the top 20% of the ranked expression profile was classified as Proneural subtype (Fisher’s Exact Test p-Value < 2.2e-16 and odds ratio = 5.4).

Other L-R pairs were specifically related to the development of the nervous system and promoting neuronal adhesion, i.e. RGMB-NEO1 and SEMA5A-PLXNB3 bindings (both in 11 out of 30 patients). The most shared interaction detected was CNTN2-NRCAM (15 out of 30 patients). The Contactin 2 (CNTN2) gene, also known as Axonal Glycoprotein TAG-1 (TAX-1), is a cell adhesion molecule that plays an important role in axonal elongation, axonal guidance, and neuronal cell migration (Masuda, 2017). The gene is also amplified and aberrantly expressed in glioblastoma, where it is involved in neoplastic glial cell migration and tumorigenesis (Rickman et al., 2001). TAX-1 binds numerous molecules, among which the Neuronal Cell Adhesion Molecule (NRCAM) gene. In response to contactin binding, NRCAM promotes directional axonal cone growth in fetal nervous system development and mediate neurite outgrowth in the peripheral nervous system (Lustig et al., 1999). Although less studied, also NRCAM is overexpressed in high-grade astrocytoma and glioblastoma, representing a marker for brain tumor detection and a putative therapeutic target (Sehgal et al., 1998).

### Cross-talk involving T-cells in glioma

The scTHI classification of non-tumor cells identified subpopulations of CD8 T cells infiltrating the microenvironment of a small subset of patients (n = 8). Although T cells play a fundamental role in antitumor immunity, glioblastoma is particularly adept at sabotaging immune surveillance causing severe T cell dysfunction, both qualitative and quantitative (Woroniecka et al., 2018). To better understand the complex role of T cells in glioma microenvironment, we simultaneously evaluated the cross-talk existing between CD8 T cells and tumor and myeloid cells, respectively.

We first investigated the putative L-R interactions occurring between tumor and CD8 T cells. Altogether, we detected 16 paracrine and 120 autocrine significant L-R pairs (**Table S8**), considering only interactions occurring in more than three patients. Most of the identified interactions involved genes of the major histocompatibility complex Class I, chemokine, interleukins, interferon gamma and Tumor Necrosis Factor (TNF) signaling genes. Among all interaction detected in paracrine mode (**Figure S6**), CADM1-CRTAM was the most common and high-scored L-R pair. The CADM1 gene, also known as TSLC1, was originally identified as a non–small-cell lung cancer tumor suppressor. The gene encodes a cell-surface protein, called Necl-2, which mediates epithelial cell junctions. We found that CRTAM receptor is highly expressed in CD8 cells with respect to other non-tumor cells in 6 out of 8 glioma patients (**Figure S7**). Although recent studies have shown a CADM1 loss at the protein and mRNA levels in high-grade (WHO III/IV) glioma compared to low-grade glioma (Houshmandi et al., 2006), the fact that we found this interaction mainly in GBM patients in association with infiltrating CD8 T cells further supports the tumor suppressor role of this gene.

Similarly, we analyzed the interactions between T CD8 and myeloid cells, identifying 53 paracrine and 159 autocrine L-R pairs (**Table S9**). The detected interactions mainly involved: (i) chemoattractant chemokine ligands and receptors, like as CXCL16/CXCR6, CXCL12/CXCR3, CLL8/CCR2, CCL5/CCR1, and others.; (ii) immune checkpoint genes, like as CD86, CD28, CTLA-4, LGALS9; (iii) lymphotoxins (LTA and LTB) and other Tumor Necrosis Factor family members, which could modulate T cells immunity through different signaling pathways. The chemokine receptor CXCR6, typically expressed on different T cell compartments, and its ligand CXCL16 secreted by macrophages cells, were found highly expressed in all patients. They mediate pro-tumorigenic effects of inflammation through direct effects on cancer cell growth and by inducing the migration and proliferation of tumor-associated leukocytes (Darash-Yahana et al., 2009). The CXCR3 receptor is also highly expressed on T cells and plays an important role in T cell trafficking and function (Groom and Luster, 2011). Lymphotoxins alfa and beta (LTA and LTB) are cytokines produced by lymphocytes, belonging to the tumor necrosis factor family. Although they were originally identified as lymphocyte products capable of exerting cytotoxic effects on tumor cells in vitro and in vivo, recent studies have shown that LT contributes to several effector responses of both the innate and adaptive immune systems. Binding to both TNFRSF1A/TNFRSF1B and LTβR with high affinity, LT-mediated signaling is essential for the development of secondary lymphoid tissues (Upadhyay and Fu, 2013). Moreover, the activation of LTβR on macrophages by T cell-derived lymphotoxins controls proinflammatory response, inducing cross-tolerance to TLR4 and TLR9 ligands through the downregulation of proinflammatory cytokine and a negative regulation of NF-kB activation induced by TLR signaling (Wimmer et al., 2012). Finally, we also identified two opposite immune checkpoint signals in almost all patients involving CD86 (n = 8) and CTLA-4 (n = 6) to binding CD28. Usually, CD86/CD28 binding results in activation and initiation of T cell effector function. However, high levels of CTLA-4 expression on T cells, probably induced by cancer, creates competition with CD28 and results in insufficient costimulation and a loss of T-cell proliferation and function (Woroniecka et al., 2018).

## DISCUSSION

In this work, we described `scTHI` a novel computational approach to identify active ligand-receptor interactions in single cells and applied to five glioma datasets encompassing 71 patients, 45,550 malignant cells and 11,510 non-malignant cells. We presented a comprehensive map of active tumor-host interactions in glioma (**Figure S8**). We have first shown that scTHI can identify recently discovered interactions validated by CODEX (Govek et al., 2019) and then explored common ligand-receptor cross-talk in glioma. Our results confirm, using a much larger scale, that myeloid cells make up the bulk of the microenvironment in glioma and that the ratio between macrophages and microglia cell increases with more aggressive phenotypes as has been previously observed for IDH-mutant glioma (Venteicher et al., 2017). Interestingly, the use of a large patient cohort allowed us to link the specific immune compartments with glioma subtypes. Indeed, we report that the presence of macrophages and myeloid-derived suppressor cells was significantly associated with the Mesenchymal subtype, as also described in (Müller et al., 2017), whereas the Proneural subtype was significantly associated with the presence of Oligodendrocytes, cells in the Classical subtype, on the other hand, tend to associated with the presence of microglia.

Our complete map of cross-talk between tumor and myeloid cells allowed us to identify some known and some novel potential targets for promoting anticancer therapy enhancing the immune response. Although a significant number of interactions are specific to few patients, when collectively analyzed, our findings show that the members of the interactions on the malignant cells enrich common pathways from those on immune cells. The ligand and receptor expressed on cancer cells participate in pathways such as extracellular matrix remodelling, cell chemotaxis, Notch signaling, axonogenesis, and gliogenesis. Instead, receptors and ligands detected on myeloid cells modulate biological processes such as leukocyte chemotaxis and migration, cell-cell adhesion, cytokine production, myeloid cell differentiation, reactive oxygen species metabolic processes and regulation of vasculature development. This pattern underlines a highly connected tumor-host communication network in glioma, where many ligands-receptors can interact on the same cell type.

We have also identified a subset of interactions that are highly conserved across different patients and datasets in paracrine mode showing that TLR2 is specifically and exclusively upregulated in glioma-associated microglia; in contrast, astrocytes and glioma cells expressed only low levels of TLR2. It is known that versican is a glioma-derived endogenous TLR2 mediator that regulates microglial MT1-MMP expression for tumor expansion (Kim et al., 2009). Microglial upregulation can be abolished by targeting TLR2 with potential therapeutic benefits in glioma progression (Hu et al., 2015). Our results confirm that TLR2 is a candidate for adjuvant therapy in the treatment of glioma (Wenger et al., 2017). We also described the interaction between EGFR and HBEGF, the feedback loop between these two genes regulates astrocytes maturation (Li et al., 2019) and promotes gliomagenesis in specific contexts, the silencing of one of the partners tends to reduce tumor growth and increases survival *in vivo* (Shin et al., 2017). Another common interaction across several patients include the pair MIF-CD74. Although MIF is a proinflammatory cytokine, it also exerts immunosuppressive functions, influencing the M1/M2 polarization of tumor-associated macrophages (Zeiner et al., 2015). In fact, recent studies have shown that the MIF-CD74 binding activates a signaling pathway resulting in M2 shift both in microglial cells, macrophages and dendritic cells (Ghoochani et al., 2016). The Inhibition of MIF signaling on these cells restore the antitumor immune response leading to a decrease in the expression of immunosuppressive factors and a reacquired capacity in cytotoxic T cells activation (Figueiredo et al., 2018). In addition, the CD74 receptor after activation is quickly internalized and recycles, therefore it constitutes an attractive target for anticancer antibody-based treatment strategies. We reported the L-R interaction including THY1 (CD90) and ITGAM/ITGB2 (Mac-1, CD11b/CD18), involved in leukocyte recruitment in response to inflammatory signals (46% of patients). The CD90 gene is a specific surface marker highly expressed in glioma-associated mesenchymal stem cells (Zhang et al., 2018). The encoded protein drives glioma progression through SRC-dependent mechanisms increasing proliferation, migration, and adhesion (Avril et al., 2017). On the other hand, the CD11b/CD18 integrins complex is abundantly expressed on monocyte/macrophages surface, where it is involved in critical adhesive reactions including the recruitment of myeloid cells to the tumor site (Podolnikova et al., 2016). Moreover, CD11b is a negative regulator of immune suppression, representing an interesting target for cancer immunotherapy (Schmid et al., 2018). In fact, CD11b activation promotes pro-inflammatory macrophage polarization, while its inhibition leads to immune suppressive macrophage polarization, vascular maturation, and accelerated tumor growth.

When we applied our algorithm in autocrine mode, we found some common interactions shared by all patients of our cohort. RPS19, for example, is an abundant intracellular protein that is expressed by virtually all cells in the body, and its extracellular functions, including interaction with C5aR1, are activated upon its release from dying cells (Yamamoto, 2007). The importance of C5aR1-RPS19 interaction for immunosuppression was recently demonstrated showing that the downregulation of RPS19 in tumor cells or pharmacological blockade of C5aR1 by C5aRA reduced this immunosuppression and led to the generation of tumor-specific T cell responses and slower tumor growth in breast and ovarian cancer cells (Markiewski et al., 2017). We have shown that the C5aR1-RPS19 interaction is among the most common signaling cross-talk between tumor cells and their microenvironment across multiple patients making these molecules an interesting target for therapeutic strategies.

Our analysis confirmed that proneural GBMs are significantly infiltrated by oligodendrocytes. MDK resulted in one of the top possible mediators of the interaction between glioma cells and oligodendrocytes through its ligand LRP2. The oncogenic role of MDK, promoting proliferation and pharmacological resistance in glioma, may involve the release of the Sonic Hedgehog (SHH) from LRP2 sequestration in oligodendrocytes (Christ et al., 2012), thus functioning to activate one of the best-known activator of proliferation and stemness of brain tumors (Niyaz et al., 2019). The single-cell characterization of proneural glioma evidenced that the concurrent expression of high levels of the PDGFA ligand by the abundant oligodendrocytes infiltrating PDGFRA-amplified proneural GBM provide the crucial initiating signaling event for the effects that this pathway has for proliferation, stemness, and progression of brain tumors (Martinho et al., 2009).

Recent trials have shown that endogenous T cells play a significant role in the prolonged survival time of glioma patients (Brown et al., 2016; Zhang et al., 2019). However, glioblastoma-induced immune suppression is a major obstacle to an effective and durable immune-mediated antitumor response. We have shown that a significant number of glioma patients present subpopulations of CD8 T cells. The presence and T cell clonal diversity in the tumor microenvironment has also been associated with response to immune-therapy in glioma (Zhao et al., 2019). Our analysis reported that this mechanism of tumor suppression could be mediated by CADM1 and its receptor CRTAM. Typically, CADM1 performs its antitumor activity ensuring that cells grow in organized layers, inhibiting uncontrolled growth. Moreover, Necl-2 binds natural killer and CD8+ T cells through a receptor known as class I-restricted T-cell-associated molecule (CRTAM), which is expressed only on activated cells (Houshmandi et al., 2006). The interaction among CRTAM and Necl-2 promotes cytotoxicity of NK cells and interferon gamma (IFN-gamma) secretion of CD8+ T cells in vitro (Boles et al., 2005).

## AVAILABILITY

scTHI is implemented in R and is available on GitHub: https://github.com/miccec/scTHI

## ACKNOWLEDGEMENTS

The research leading to these results has received funding from Associazione Italiana per la Ricerca sul Cancro (AIRC) under IG 2018 - ID. 21846 project.

## STAR METHODS

### Single Cell Datasets

Glioma single cell gene expression profiles were collected from six different dataset of glioma. We obtained a subset of IDH mutant gliomas including 10,688 cells from 16 patients from two different studies (Tirosh et al., 2016a; Venteicher et al., 2017). Later, we will refer to this subset of cell as a unique dataset. High grade glioma profiles were collected from three distinct studies: Darmanis et al. profiled 3,589 cells from 4 patients (Darmanis et al., 2017); Yuan et al. profiled ∼ 29,000 cells from 10 patients augmented with two novel specimens (PJ052 and PJ053) following library construction and sequencing described in (Yuan et al., 2018); Neftel et al. profiled 7,930 cells from 28 patients (Neftel et al., 2019). We also considered another dataset of 5,603 single cell profiles derived from both LGG and HGG patients (n = 13) (Yu et al., 2019). Overall, we collected gene expression profiles of 57,060 cells for a total of 71 glioma patients. The cohort is composed of several tumor histologies, including 7 Oligodendroglioma (4,753 cells), 2 Oligodendroastrocytoma (612 cells), 11 Astrocytoma (9,421 cells) and 50 Glioblastoma (42,274 cells). Gene expression profiles was processed independently for each dataset. The TPM normalized data of the Tirosh et al., Venteicher et al. and Neftel et al. (Neftel et al., 2019; Tirosh et al., 2016a; Venteicher et al., 2017) datasets were downloaded from the GEO repositories under accession numbers GSE70630, GSE89567, GSE131928, respectively. Gene expression profiles from Darmanis et al. and Yuan et al. (Darmanis et al., 2017; Yuan et al., 2018) were downloaded from GEO repositories under accession numbers GSE84465 and GSE103224, respectively. While, raw data from Yu et al. (Yu et al., 2019) were obtained from authors. The last three datasets were further processed applying a library size normalization and logarithmic transformation. Moreover, in order to reduce the drop-out effects, data matrices were also imputed with a Markov Affinity-based graph approach (van Dijk et al., 2018). When the specific information to distinguish malignant cells from non-tumor cells was not available, we analyzed the chromosomal aberrations in each individual cell by averaging expression level along genomic locations as performed by InferCNV (Patel et al., 2014). The chromosomal landscape of inferred CNV allows us to identify non-transformed cells, i.e. cells that didn’t harbor the typical chromosomal alterations observed in glioma. Altogether we identified 45,550 malignant cells and 11,510 non-malignant cells among datasets.

### Ligand-receptor collection

In order to identify potential tumor-host interactions we collected a list of 2,548 pairs of ligands and receptors (**Table S1**) curating publicly available resources (Ramilowski et al., 2015; Vento-Tormo et al., 2018). The curated list is composed of known and novel literature-supported interactions and includes both heteromeric and monomeric ligands/receptors mainly related to chemokine, cytokine, growth factors, integrin, TGF and TNF family members, semaphorins, ephrins, Wnt and Notch signalings. The table of interactions is released in the `scTHI` tool on github.

### Immune cell type classification

Cell-type specific signatures and markers were used to infer the cellular identity of non-malignant cell subpopulations. For this purpose, we generated a curated list of 295 signatures of the Immune and Central Nervous Systems integrating data from various sources (**Table S2**). In particular we merged a manual collection of marker genes with a set of signatures available from public databases and published studies, including: (i) a compendium of 64 human cell types signatures including lymphoid, myeloid, stromal, tissue-specific and stem cells, collected from FANTOM5, ENCODE, Blueprint and Gene Expression Omnibus (GEO) data portals (Aran et al., 2017); (ii) a set of markers for 30 immune cell types, including myeloid and lymphoid subpopulations identified from Peripheral Blood Mononuclear Cells (PBMCs) (Butler et al., 2018); (iii) a set of Central Nervous System cell signatures including astrocytes, neuron, oligodendrocytes, microglia, and endothelial cells (Zhang et al., 2014); (iv) a set of 53 signatures corresponding to 26 different cell types (Bindea et al., 2013; Charoentong et al., 2017; Rooney et al., 2015; Tirosh et al., 2016b); (v) two gene expression programs related to microglia and bone marrow-derived macrophages in gliomas (Venteicher et al., 2017). All the 295 signatures are released in the `scTHI` tool.

To evaluate the enrichment of each immune cell type in glioma samples we used Normalized Enrichment Score (NES) of the Mann Whitney Wilcoxon Gene Set test (mww-GST) that we previously described in (Frattini et al., 2018). Briefly, NES is an estimate of the probability that the expression of a gene in the gene set is greater than the expression of a gene outside this set:

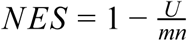

where *m* is the number of genes in a gene set, *n* is the number of those outside the gene set, *U* = *mn* + *m*(*m* + 1) − *T*, and *T* is the sum of the ranks of the genes in the gene set. We applied single cell mww-GST and classified each cell according to the signature with the highest NES and corrected p-value less than 0.01.

### scTHI scores

The scTHI scores are computed between pairs of clusters of single cells assigning a score to each interaction of the table. A typical example is when we have one cluster from the microenvironment (ex. macrophages) and one cluster from the tumor cells. Given a single cell profile *G*, ranked from the high expressed to the low expressed genes, we call *G*^*T*^ the set of genes in the top of the ranked profile (in all the reported experiments we use the top 20%, the accompanying code allows to select the threshold). Let also call the two cluster *A* and *B*, then for every ligand-receptor (L-R) pair of the interaction table we compute the following score:

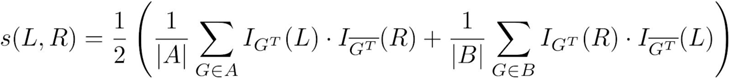

where *I* is the indicator function

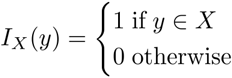

Briefly, the score *s* is the average between two percentages: the percentage of cells in cluster *A* where the gene *L* is at the top of the ranked list and gene *R* is not at the top and vice versa for cluster *B* in the second percentage, it tends to give a higher score to paracrine interactions (in order to also score autocrine interactions the second term in the product of the two summations in equation can be removed). A null distribution of the interaction score to assign significance is then obtained by a bootstrap procedure shuffling the input data.

### Gene Ontology enrichment analysis

The GO category enrichments of ligand and receptor genes detected by `scTHI` analysis were performed using clusterProfiler (Yu et al., 2012). Enriched GO terms were filtered using an adjusted p-Value cutoff of 0.0001.

## TABLE AND FIGURES LEGENDS

**Table S1. Ligand-Receptor interactions.** List of Ligand-Receptor interaction pairs provided in `scTHI`.

**Table S2. Signatures.** List of immune system and tissue cell type signatures provided in `scTHI.`

**Table S3. Non tumor cell classification.** Phenotype classification of tumor microenvironment cells in glioma datasets performed by TME.classification function provided by `scTHI`.

**Table S4. Tumor-Myeloid cell interactions.** List of significant paracrine and autocrine L-R interactions occurring between tumor and myeloid cells from considered datasets detected by `scTHI.`

**Table S5. Enriched GO categories of L-R partners in tumor and myeloid cells.** List of significant enriched GO biological processes of ligand and receptor genes expressed in tumor and myeloid cells, respectively.

**Table S6. Tumor-Oligodendrocyte cell interactions.** List of significant paracrine and autocrine L-R interactions occurring between tumor and oligodendrocyte cells from considered datasets detected by `scTHI.`

**Table S7. Enriched GO categories of L-R partners in tumor and oligodendrocyte cells.** List of significant enriched GO biological processes of ligand and receptor genes expressed in tumor and oligodendrocyte cells, respectively.

**Table S8. Tumor-CD8 cell interactions.** List of significant paracrine and autocrine L-R interactions occurring between tumor and CD8 T cells detected by `scTHI.`

**Table S9. Myeloid-CD8 cell interactions.** List of significant paracrine and autocrine L-R interactions occurring between myeloid and CD8 T cells detected by `scTHI.`

---

**Figure S1.**
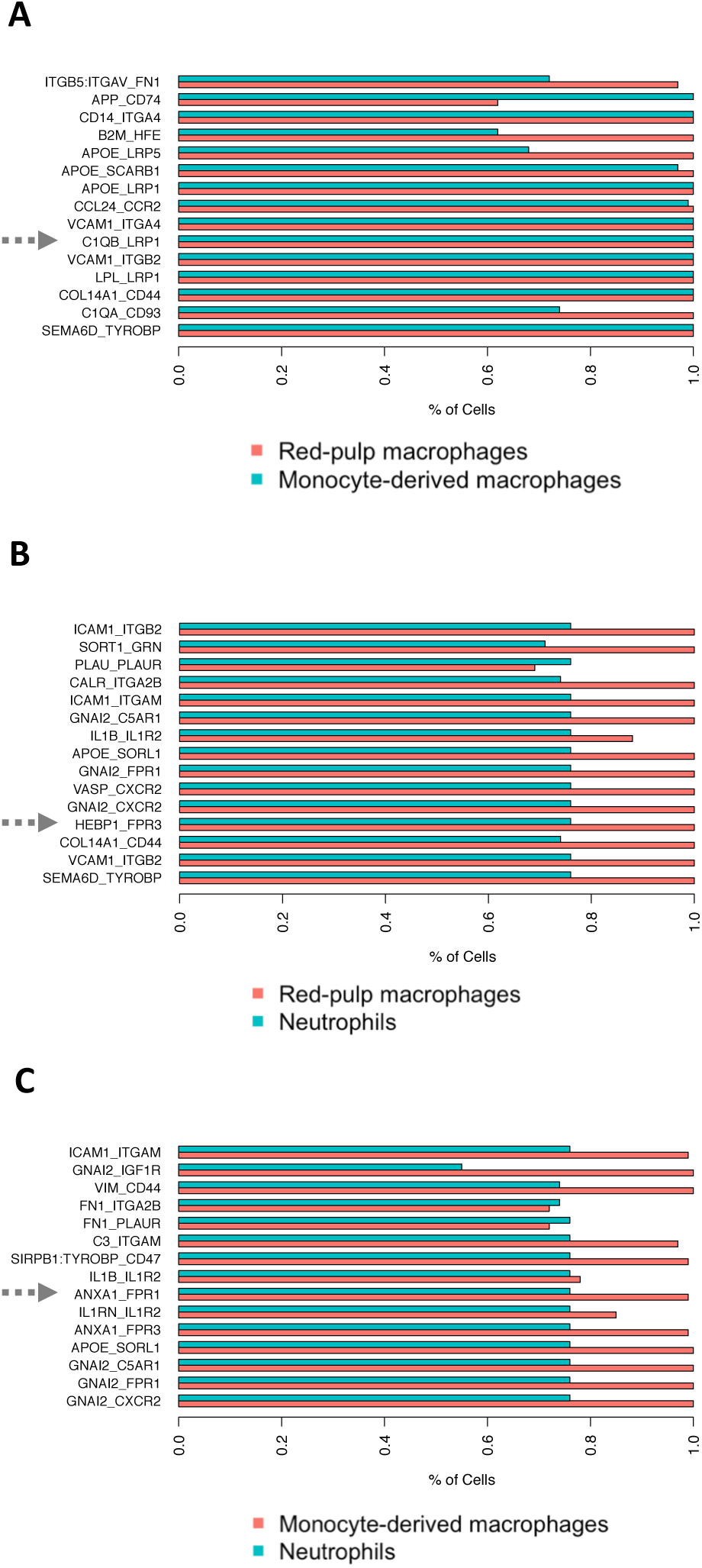
scTHI recoves validated interactions. The barplots show the top 15 significant interactions identified between (A) red-pulp and monocyte-derived macrophage cells; (B) red-pulp macrophages and neutrophils; (C) monocyte-derived macrophages and neutrophils. The interactions spatially resolved in ((15) are pointed out with an arrow.

**Figure S2.**
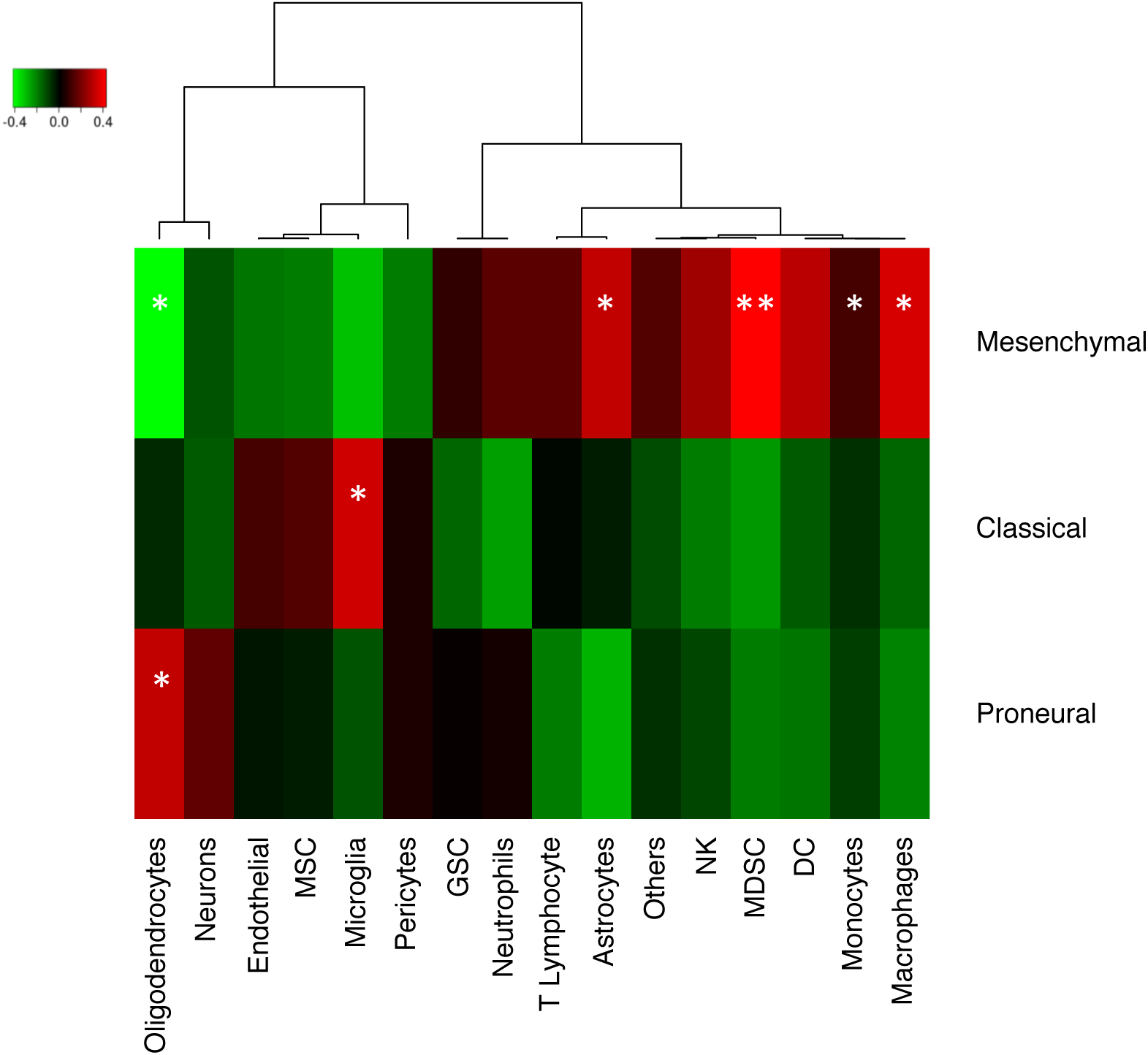
Association between immune cell types and glioma subtypes. Heatmap of correlation between the percentage of cells classified in one of the glioma subtypes with the percentages of non tumor cell types. Significant correlations are marker with * (p-value <= 0.05) and ** (p-value <= 0.01).

**Figure S3.**
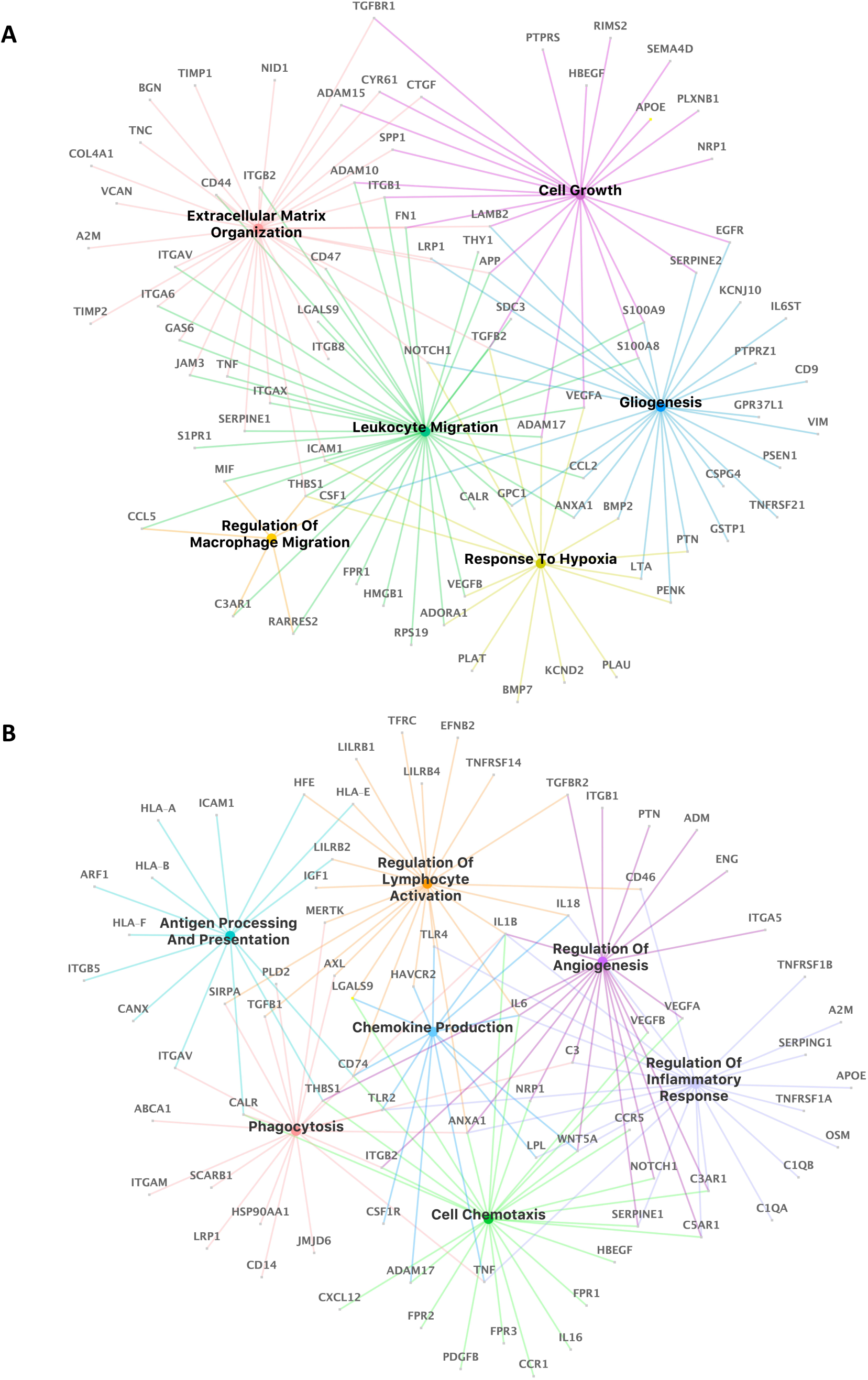
Enriched GO categories of L-R pair. Network of most represented biological process category enriched by L-R partners on tumor (A) and on myeloid cells (B).

**Figure S4.**
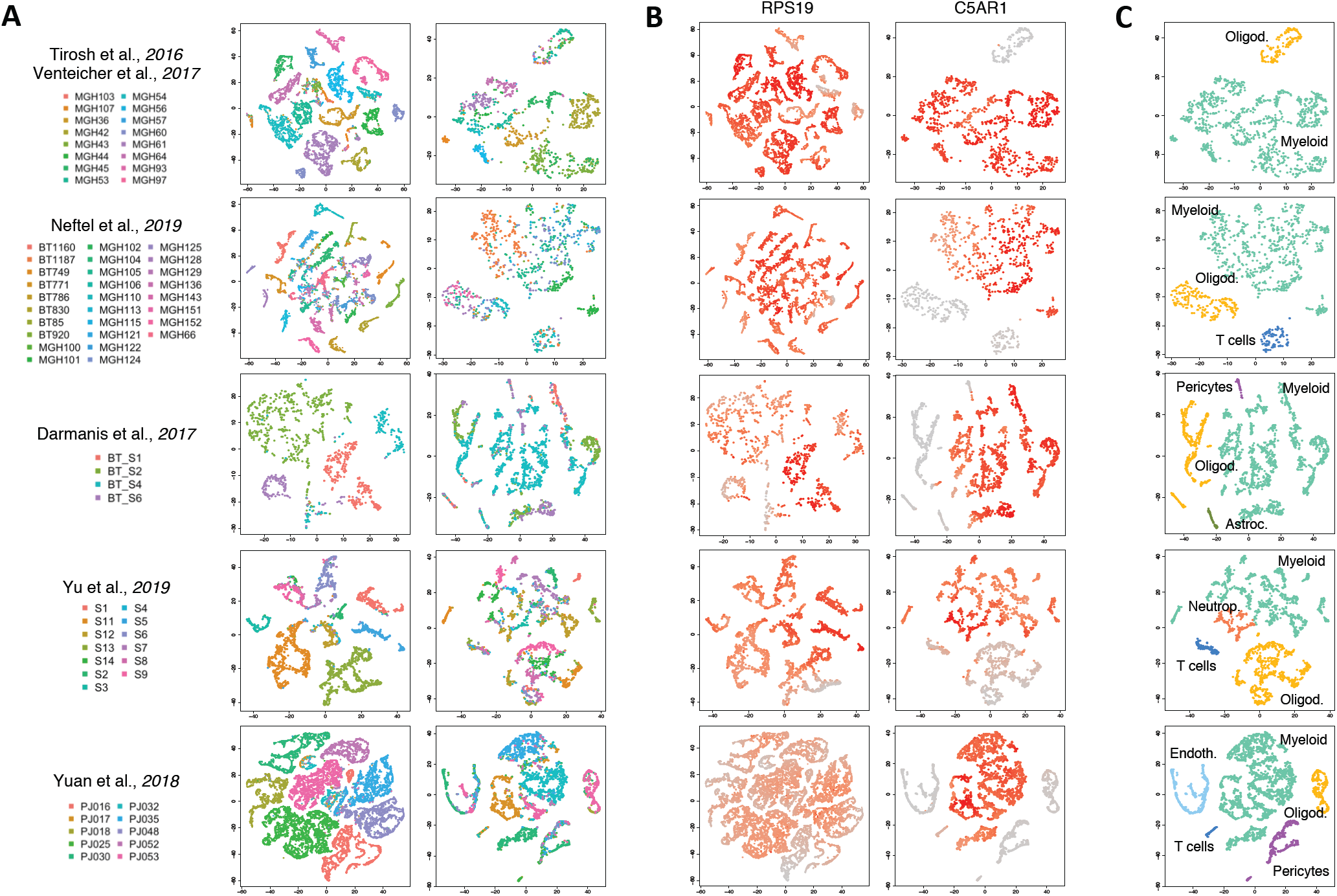
RPS19-C5AR1 interaction. TSNE plots of tumor and non-tumor cells for each dataset analyzed. (A) Each cell is colored by patient (on the right non-tumor and on the left tumor cells). (B) Each cell is colored according to expression value of the genes RPS19 and C5AR1 in tumor and non tumor cells, respectively. (C) tSNE plot of non-tumor cells colored according to cell type classification.

**Figure S5.**
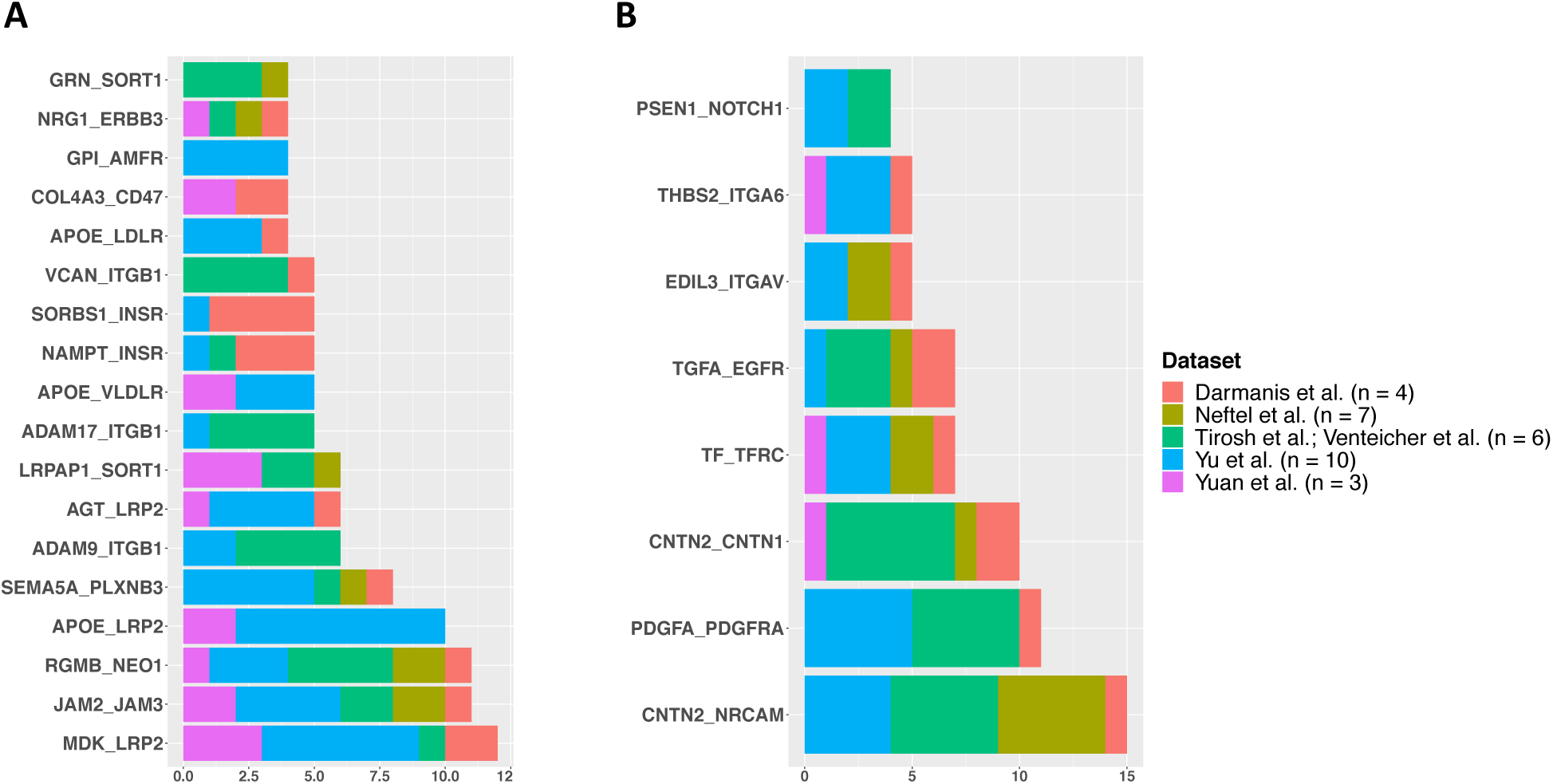
Paracrine tumor-oligodendrocyte cell interactions. Barplots show significant paracrine L-R interactions (p-Value <= 0.05 and scTHI score >= 0.50) occurring between tumor and oligodendrocyte cells shared in at least 4 patients. On the x axis are shown the number of patients where each interaction occurred. (A) Interaction pairs in which the ligand is expressed on tumor cells and the receptor on oligodendrocyte cells. (B) Interaction pairs in which the ligand is expressed on oligodendrocyte cells and the receptor on tumor cells.

**Figure S6.**
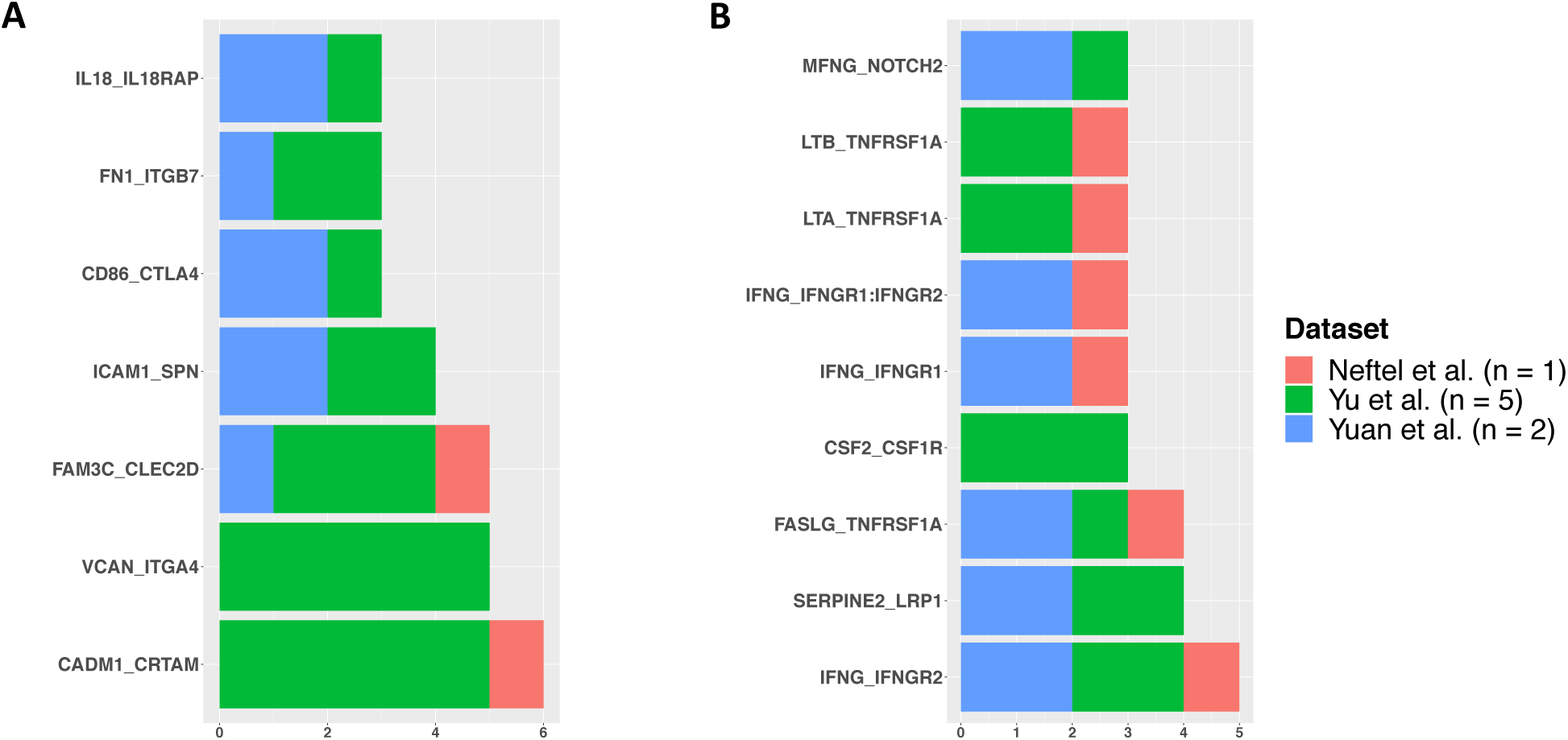
Paracrine tumor-CD8 cell interactions. Barplots show significant paracrine L-R interactions (p-Value <= 0.05 and scTHI score >= 0.50) occurring between tumor and CD8 cells shared in at least 4 patients. On the x axis the number of patients where each interaction occurred is shown. (A) Interaction pairs in which the ligand is expressed on tumor cells and the receptor on CD8 cells. (B) Interaction pairs in which the ligand is expressed on CD8 cells and the receptor on tumor cells.

**Figure S7.**
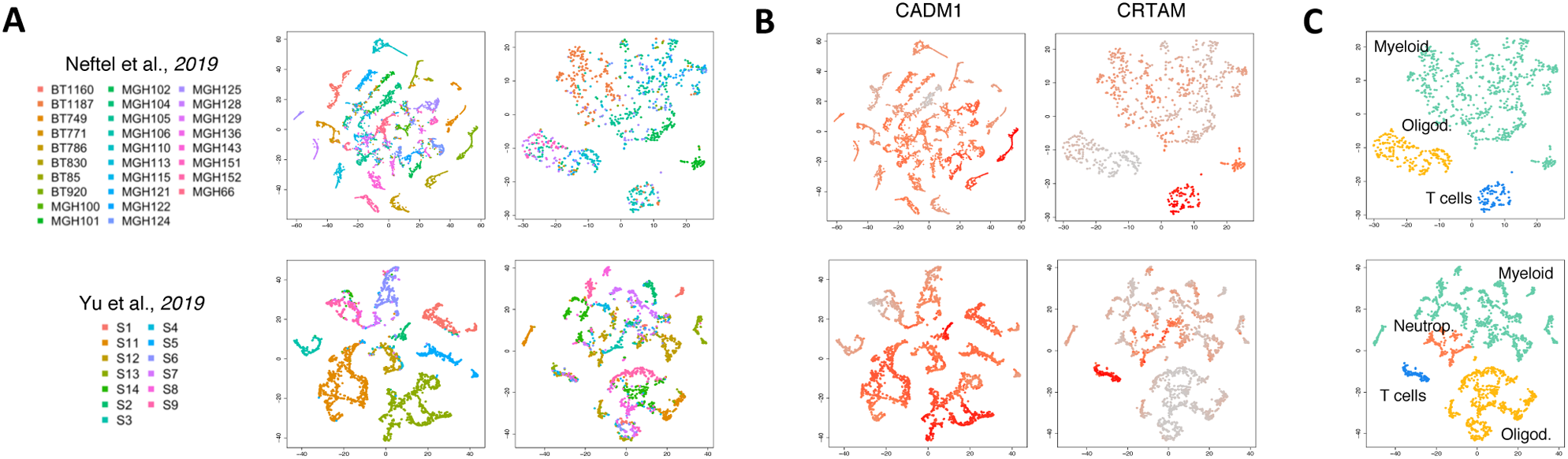
CADM1-CRTAM interaction. TSNE plots of tumor and non-tumor cells for each dataset analyzed. (A) Each cell is colored by patient (on the right non-tumor and on the left tumor cells). (B) Each cell is colored according to expression value of the genes CADM1 and CRTAM in tumor and non tumor cells, respectively. (C) tSNE plot of non-tumor cells colored according to cell type classification.

**Figure S8.**
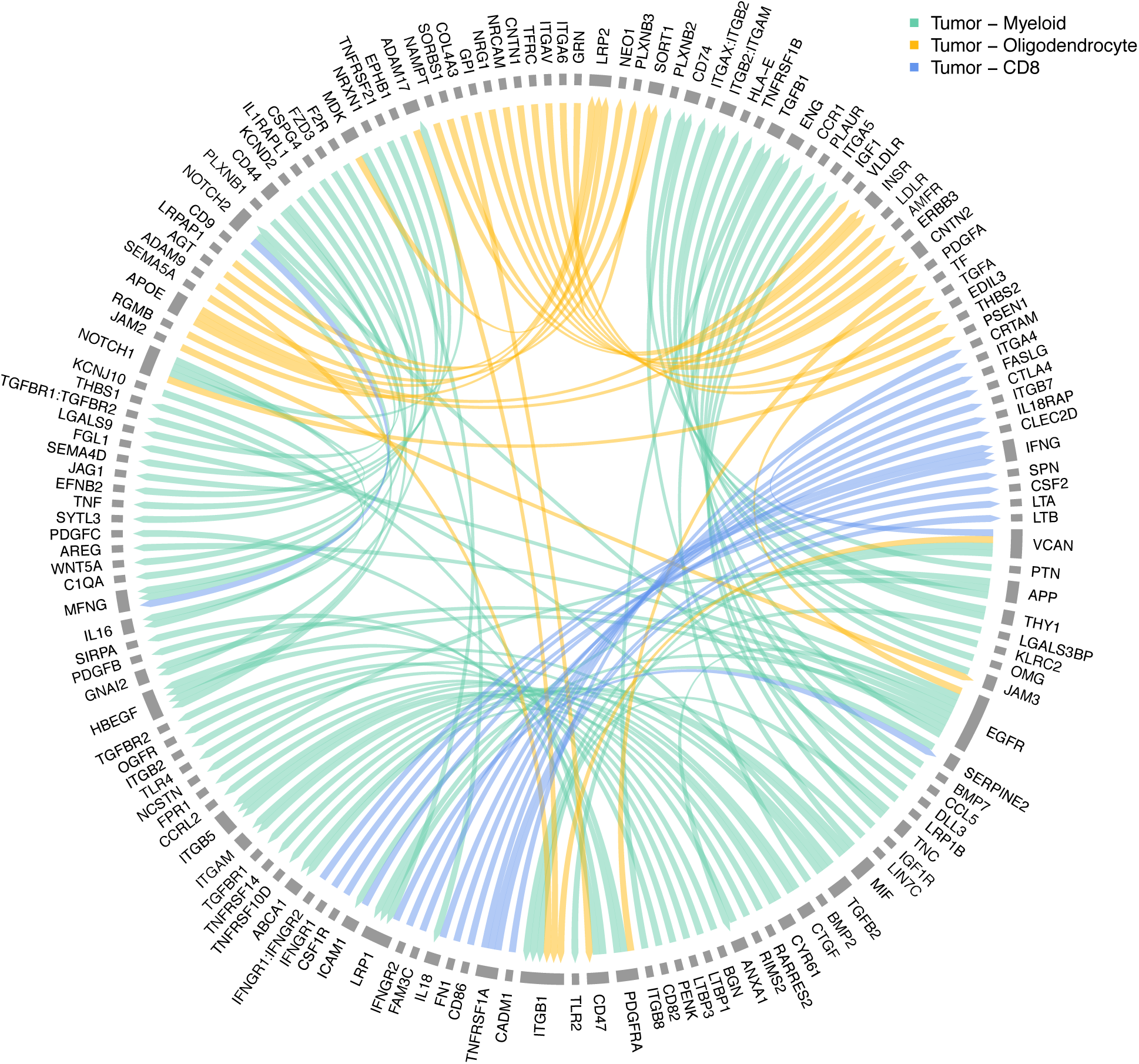
A map of tumor-host interactions in glioma. Chord diagram of paracrine tumor-host interaction detected by scTHI. The color of the arcs indicates the clusters of cells among which the interaction has been identified. The origin of the arch indicates that the ligand or receptor is expressed on the tumor, instead the arrow of the arch indicates that the ligand or receptor is expressed on the cells of the microenvironment.

